# High-Throughput Empirical and Virtual Screening to Discover Novel Inhibitors of Polyploid Giant Cancer Cells in Breast Cancer

**DOI:** 10.1101/2024.09.23.614522

**Authors:** Yushu Ma, Chien-Hung Shih, Jinxiong Cheng, Hsiao-Chun Chen, Li-Ju Wang, Yanhao Tan, Yu-Chiao Chiu, Yu-Chih Chen

**Affiliations:** UPMC Hillman Cancer Center, University of Pittsburgh, 5115 Centre Ave, Pittsburgh, PA 15232, USA; Department of Computational and Systems Biology, University of Pittsburgh, 3420 Forbes Avenue, Pittsburgh, PA 15260, USA; Department of Bioengineering, Swanson School of Engineering, University of Pittsburgh, 3700 O’Hara Street, Pittsburgh, PA 15260, USA; Division of Malignant Hematology and Medical Oncology, Department of Medicine, University of Pittsburgh, 5150 Centre Avenue, Pittsburgh, PA 15232, USA; CMU-Pitt Ph.D. Program in Computational Biology, University of Pittsburgh, 3420 Forbes Avenue, Pittsburgh, PA 15260, USA

**Keywords:** Polyploid Giant Cancer Cells, Machine Learning, Single-Cell Analysis, Treatment Resistance, Breast Cancer

## Abstract

Therapy resistance in breast cancer is increasingly attributed to polyploid giant cancer cells (PGCCs), which arise through whole-genome doubling and exhibit heightened resilience to standard treatments. Characterized by enlarged nuclei and increased DNA content, these cells tend to be dormant under therapeutic stress, driving disease relapse. Despite their critical role in resistance, strategies to effectively target PGCCs are limited, largely due to the lack of high-throughput methods for assessing their viability. Traditional assays lack the sensitivity needed to detect PGCC-specific elimination, prompting the development of novel approaches. To address this challenge, we developed a high-throughput single-cell morphological analysis workflow designed to differentiate compounds that selectively inhibit non-PGCCs, PGCCs, or both. Using this method, we screened a library of 2,726 FDA Phase 1-approved drugs, identifying promising anti-PGCC candidates, including proteasome inhibitors, FOXM1, CHK, and macrocyclic lactones. Notably, RNA-Seq analysis of cells treated with the macrocyclic lactone Pyronaridine revealed AXL inhibition as a potential strategy for targeting PGCCs. Although our single-cell morphological analysis pipeline is powerful, empirically testing all existing compounds is impractical and inefficient. To overcome this limitation, we trained a machine learning model to predict anti-PGCC efficacy *in silico*, integrating chemical fingerprints and compound descriptions from prior publications and databases. The model demonstrated a high correlation with experimental outcomes and predicted efficacious compounds in an expanded library of over 6,000 drugs. Among the top-ranked predictions, we experimentally validated two compounds as potent PGCC inhibitors. These findings underscore the synergistic potential of integrating high-throughput empirical screening with machine learning-based virtual screening to accelerate the discovery of novel therapies, particularly for targeting therapy-resistant PGCCs in breast cancer.

## Introduction

PGCCs are cancer cells with additional copies of chromosomes, often resulting in significantly larger cell size and increased genomic content.^1–3^ These cells are found across various cancer types, including breast, prostate, lung, ovarian and colorectal cancers.^4–8^ The presence of PGCCs has been correlated with advanced disease stages, increased tumor aggressiveness, and poor clinical outcomes. The formation of PGCCs can be attributed to several mechanisms, including aberrant cell cycle regulation, mitotic failure, and response to cellular stress such as chemotherapy and radiation. These mechanisms result in the cells bypassing normal mitotic checkpoints, leading to endoreduplication or cell fusion events that contribute to polyploidy.^9–15^ PGCCs contribute significantly to tumor heterogeneity. By re-shuffling genomic content of multiple copies of genome,^16^ they generate diverse progeny through asymmetric division and budding allows for the rapid adaptation of tumor cells to changing microenvironments and therapeutic pressures.^17^ This adaptability promotes tumor evolution and metastasis, complicating treatment strategies.

PGCCs have emerged as a key target in cancer research due to their critical role in therapy resistance. These cells exhibit resistance to conventional chemotherapies and radiation therapy, often surviving initial treatments and giving rise to recurrent tumors.^15, 18, 19^ This resistance is mediated through multiple mechanisms, including enhanced DNA repair capabilities, activation of survival pathways, avoidance of apoptosis, and the ability to enter a dormant state. In addition, PGCCs are reported to exhibit stem cell-like properties by their enhanced tumor-initiating capability and up-regulation of relevant biomarkers.^20–22^ Their presence often correlates with more aggressive disease phenotypes and poorer patient outcomes. Targeting PGCCs represents a promising therapeutic strategy. Approaches under investigation include disrupting the specific cell cycle and survival pathways active in PGCCs, as well as exploiting their unique metabolic dependencies.^23–28^ Therapies aimed at eliminating PGCCs or preventing their formation could enhance treatment efficacy and reduce relapse rates.

Although there has been some progress in this direction, to date,^23–32^ there are no effective therapies targeting PGCCs.^15^ The development of anti-PGCC treatments has been hindered by the absence of a high-throughput method to rapidly quantify these cells. Traditional drug screening assays, such as MTT, XTT, or ATP, quickly measure the overall inhibition of cancer cell populations but fail to provide specific information on the elimination of a small PGCC subpopulation, which are crucial for addressing treatment resistance and relapse. PGCCs can be characterized by the excessive DNA content and large cell and nuclear size. Currently, the gold standard for identifying and isolating PGCCs involves fluorescence-activated cell sorting (FACS) combined with visual confirmation.^20^ While flow cytometry can quantify the number and percentage of PGCCs, it is impractical for screening thousands of compounds or for monitoring the dynamic processes of PGCC induction and death. The limitations of existing approaches underscore the need for a high-throughput and precise analytical method specifically tailored for PGCC research. Building upon the advances in image-based cell segmentation and detection methods,^33–37^ we recently established a dedicated workflow for the identification and tracking of PGCCs.^38^ In this study, we expanded the screen to a library of 2,726 FDA Phase 1-approved drugs to identify novel PGCC inhibitors. Additionally, we conducted RNA-Seq analysis to preliminarily elucidate the mechanisms of these anti-PGCC compounds and to explore new strategies for targeting PGCCs.

Although our single-cell morphological analysis allows high-throughput testing of thousands of compounds, it is impractical to empirically test all existing compounds. This challenge underscores the need for computational methods that can efficiently predict anti-PGCC drug responses, streamlining the drug discovery process by identifying promising candidates for experimental validation. Machine learning models have emerged as powerful tools, offering a promising solution by leveraging multi-omics data and biochemical features of compounds, such as chemical structures, to predict drug sensitivity across cancer cell lines.^39–45^ However, to the best of our knowledge, no machine learning models currently exist for predicting anti-PGCC compounds, largely due to the lack of large training datasets. Establishing such methods is essential for advancing the development of targeted therapies against these challenging cancer cells. In this study, powered by our high-throughput morphological assay, we systematically evaluated a wide array of machine learning models to predict anti-PGCC effects (**Fig. 1a**). Furthermore, we developed a novel ensemble model that integrates biochemical features with pharmacological descriptions of compounds to enhance prediction performance. This model enabled virtual screening of an expanded library of 6,575 compounds for potential drug repurposing opportunities. Among the top predictions, we experimentally validated two compounds. Taken together, this study demonstrates the significant potential of integrating empirical and virtual screening approaches for PGCCs, which may unlock new avenues for overcoming cancer therapy resistance and ultimately lead to improved patient outcomes.

**Figure 1.**
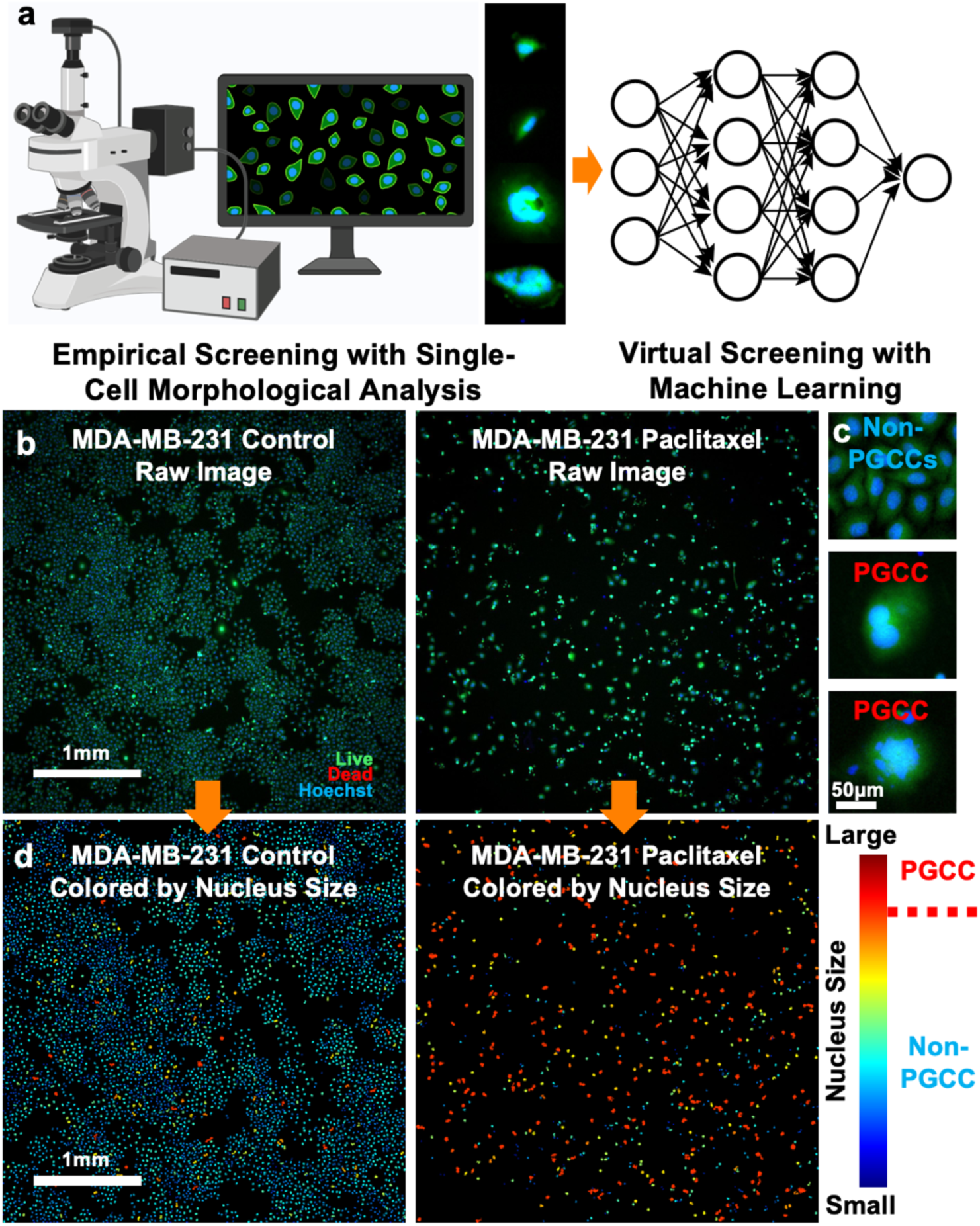
Single-cell morphological analysis for PGCC identification. (a) A conceptual diagram illustrating empirical drug screening by single-cell morphological analysis and virtual screening by machine learning. (b) Raw fluorescent images of MDA-MB-231 cells treated with or without 10 µM Paclitaxel (Scale bar: 1 mm). Cells were stained with Live (green), Dead (red), and Hoechst (blue) reagents. (c) Enlarged images of representative MDA-MB-231 PGCCs and non-PGCCs (Scale bar: 50 μm). (d) Our single-cell morphological analysis pipeline converts raw images to pseudo-colors indicating nuclear size: red for larger nuclei and blue for smaller nuclei.

## Methods

### Cell culture

We cultured MDA-MB-231 and Vari068 cells in Dulbecco’s Modified Eagle Medium (DMEM, Gibco 11995) supplemented with 10% fetal bovine serum (FBS, Gibco 16000), 1% GlutaMax (Gibco 35050), 1% penicillin/streptomycin (pen/strep, Gibco 15070), and 0.1% plasmocin (InvivoGen ant-mpp). SUM159 cells were cultured in F-12 medium (Gibco 11765) supplemented with 5% FBS (Gibco 16000), 1% pen/strep (Gibco 15070), 1% GlutaMax (Gibco 35050), 1 μg/mL hydrocortisone (Sigma H4001), 5 μg/mL insulin (Sigma I6634), and 0.1% Plasmocin (InvivoGen ant-mpp). MDA-MB-231 and SUM159 cells were obtained from Dr. Gary Luker’s lab at the University of Michigan, while Vari068 cells were obtained from Dr. Max Wicha’s lab at the University of Michigan. The Vari068 cells, derived from an ER-/PR-/Her2-breast cancer patient who provided informed consent, were adapted to a standard two-dimensional culture environment.^46–48^ All cell cultures were maintained at 37 °C in a humidified incubator with 5% CO2 and passaged upon reaching over 80% confluency. All cell lines were cultured with a mycoplasma antibiotic Plasmocin.

### Compound screening to identify inhibitors of PGCCs

In our screening experiments, we utilized a compound library of 2,726 compounds, each having successfully completed Phase I drug safety confirmation (APExBIO, L1052, DiscoveryProbe™ Clinical & FDA Approved Drug Library). These compounds were prepared at a concentration of 10 mM in DMSO or PBS. For screening, serial dilution was performed to achieve a final concentration of 10 µM. DMSO at 0.1% was used as the control treatment. Cells were harvested from culture dishes using 0.05% Trypsin/EDTA (Gibco, 25200), centrifuged at 1,000 rpm for 4 minutes, re-suspended in appropriate media, and seeded into 96-well plates. The number of cells seeded per well varied by cell line: 1,000 for SUM159 and MDA-MB-231 in 100 μL of media per well. Cells were cultured for 24 hours before treatment with compounds for 48 hours. Post-treatment, cells were stained with 0.3 μM Calcein AM (Biotium, 80011-2), 0.6 μM Ethidium homodimer-1 (Invitrogen™, L3224 Live/Dead Viability/Cytotoxicity Kit), and 8 μM Hoechst 33342 (Thermo Scientific 62249), followed by a 30-minute incubation. For other experiments, 4,000 cells per well were seeded for all cell lines. After 24 hours, cells were treated with PGCC-inducing agents (Docetaxel 1 μM) for 48 hours. Post-induction, the reagents were aspirated, and the test compounds were added to treat the mixed populations for an additional 48 hours without flow sorting. The same staining and imaging protocol was used to quantify PGCCs and non-PGCCs after treatments.

### Image acquisition

Cells in 96-well plates were imaged using an inverted Nikon Ti2E microscope. Brightfield and fluorescence images were captured with a 4x objective lens and a Hamamatsu ORCA-Fusion Gen-III SCMOS monochrome camera. Each field of view covers approximately 14 mm^2^, accommodating up to 10,000 cells per image. Hoechst-stained cell nuclei were visualized with a DAPI filter set, while live and dead cells were detected using FITC and TRITC filter sets, respectively. Auto-focusing ensured image clarity, with the entire imaging process for a 96-well plate completed in under 9 minutes.

### Single-cell morphological analysis software

The goal of our image processing is to quantify viable cells and distinguish PGCCs from non-PGCCs. We developed a custom MATLAB (2021b) program to achieve this in three steps: (1) identify cell nuclei with Hoechst staining, (2) determine cell viability, and (3) recognize PGCCs based on nuclear size. Hoechst-stained images were initially filtered using top-hat and bottom-hat filters to reduce background noise, then enhanced through contrast adjustment, and binarized to measure nuclear sizes. Cell debris was excluded based on smaller sizes.^49^ Live/Dead staining was employed to exclude dead cells, identified by dim Live signals and bright Dead signals. The cell counting method was adapted from our previous work.^50–52^ Live cells with nuclei larger than 300 pixels using a 4X objective lens or 1,875 pixels using a 10X objective lens (817 µm^2^ area, equivalent to a 32 µm diameter circle) were classified as PGCCs, while smaller nuclei were considered non-PGCCs. These thresholds were empirically validated with flow cytometry and visual confirmation (Fig. 1). Among the 2,726 compounds, 29 compounds were excluded due to their fluorescent colors which interfere with image processing.

### Whole-transcriptome sequencing

We extracted RNA from MDA-MB-231 cells, both untreated and treated with 1 μM Pyronaridine Tetraphosphate for 2 days, using the PureLink™ RNA Mini Kit (Invitrogen™, 12183018A). The RNA samples were processed at the UPMC Hillman Cancer Center Cancer Genomics Facility with a KAPA RNA HyperPrep Kit with RiboErase. Each sample population was expected to generate approximately 40 million reads (38×38 base paired-end), with two biological replicates conducted. Reads were aligned using Bowtie2 read aligners in Partek, followed by transcriptome assembly and differential expression analysis with DESeq2.^53, 54^

### Functional enrichment analysis of the Pyronaridine treatment

Gene Set Enrichment Analysis (GSEA) was performed to understand the underlying mechanisms of Pyronaridine treatment.^55^ Genes from RNA-seq were ranked based on the statistical significance (*P*-value) of their differential expression in Pyronaridine-treated MDA-MB-231 cells compared to untreated cells. The curated gene sets representing genetic and chemical perturbations (CGPs) from the Molecular Signatures Database (MSigDB) were tested for enrichment at the negative end of the ranked gene list (*i.e.*, downregulated genes in response to Pyronaridine).^56^ To analyze overlaps among enriched gene sets, we utilized EnrichmentMap and AutoAnnotate in Cytoscape for constructing and visualizing a gene set association network.^57^ Gene set associations were represented by the degree of gene overlap between two sets, calculated as the average of the Jaccard index and the overlap coefficient (referred to as the combined coefficient). Gene sets with an FDR *q*-value below 0.05 in GSEA and a combined coefficient above 0.375 were included in the association network. Additionally, we analyzed the leading-edge subset of an enriched gene set of interest identified by GSEA, which represents the top-ranked genes that contribute most to the enrichment score. This subset was further studied for its potential relevance in the response to Pyronaridine.

### Statistical analysis

Statistical analyses were conducted using R (version 4.1), GraphPad Prism 10, and MATLAB. GraphPad Prism 10 software determined half-maximal inhibitory concentrations (IC50s). Two-tailed Student’s *t*-test compared two groups, while paired 1-way ANOVA and Fisher’s Least Significant Difference (LSD) test compared multiple groups, considering treatment conditions as the variable. Within each cell line, treated versus untreated conditions were consistently paired for comparisons, with significance set at *P*<0.05. The standard deviation was represented by error bars; sample/group details were specified in figure captions. For data with high variability (*e.g.*, gene expression levels), comparisons were made on a log scale.

### Representation of drug features using structures and descriptions

For machine learning modeling, each drug was represented by either a vector of molecular fingerprints to capture its biochemical and structural features, or a vector of text embeddings to encode descriptions of its pharmacological, biochemical, and molecular biological properties. Drug structures were represented by the Simplified Molecular Input Line Entry System (SMILES) line notation. Canonical SMILES codes were obtained from PubChem using the Python PubChemPy package and then converted into molecular fingerprints based on the Molecular ACCess System (MACCS), PubChem, and Extended-Connectivity Fingerprint (ECFP6) systems using the R rcdk package.^58^ The molecular fingerprints are binary vectors that encode the structural properties of a drug, with lengths of 166, 881, and 1,024 bits, respectively, where each bit denotes the presence (1) or absence (0) of a pre-defined structural property. Text descriptions of drugs were obtained from PubChem using the PUG REST interface, which provides programmatic access to PubChem data.^59, 60^ We then converted the descriptions into text embeddings using the latest embedding methods developed by OpenAI, including text-embedding-3-small (1,536 dimensions) and text-embedding-3-large (3,072 dimensions), which generate vectors composed of continuous values to represent the semantic information of drug descriptions.

### Machine learning models to predict anti-PGCC efficacy

We trained machine learning models to predict drug responses in PGCCs of MDA-MB-231 based on drug structures and descriptions. The normalized count of PGCCs, compared between treated and untreated cells, was increased by 10^-3^ and then log2-transformed and used as the prediction target. We employed 10-fold cross-validations to train and test each model. In each round of 10-fold cross-validation, the drugs were randomly partitioned into 10 sets, where 9 sets were used for model training and the remaining set was used for testing by calculating the Pearson correlation coefficient between the actual and predicted values. Once all 10 sets were tested by the corresponding trained models, we summarized the performance by averaging the 10 correlation coefficients. This entire process, including random partitioning and 10-fold cross-validation, was repeated for 10 rounds. The results from these 10 rounds are presented in box plots, with performance summarized by the median correlation value. We evaluated a total of seven linear and nonlinear regression-based machine learning models, including linear regression with L2 regularization (Ridge), support vector machine (SVM), random forest (RF), histogram-based gradient boosting (HGB), decision tree (DT), stochastic gradient descent linear regression (SGD), and multi-layer perceptron (MLP). These models were implemented using the respective functions of the Python scikit-learn library. For ensemble learning, the predicted drug responses from two individual models, trained on either drug structures or descriptions, were used as inputs for training a linear regression model to predict the drug response. We ensured that all random partitions were applied consistently across individual and ensemble models to allow for rigorous comparison of the results.

## Results and Discussion

### Comprehensive compound efficacy analysis by quantifying PGCCs and non-PGCCs

We developed a single-cell morphological analysis pipeline to rapidly quantify PGCCs and non-PGCCs by identifying cell nuclei with Hoechst staining, excluding dead cells using Live/Dead staining, and distinguishing PGCCs and non-PGCCs based on nuclear size (**Fig. 1a**).^38^ This pipeline was validated with multiple breast cancer cell lines and confirmed through flow cytometry and visual inspection. As a demonstration, we treated MDA-MB-231 cells with Paclitaxel, a common and widely used drug for triple-negative breast cancer (TNBC) (**Fig. 1b**). Without treatment, the cell population was predominantly non-PGCCs and much higher in number. Paclitaxel treatment significantly reduced the total number of cells while inducing a higher proportion of PGCCs, which can be the source of treatment resistance. **Fig. 1c** shows enlarged views of non-PGCCs and PGCCs. Our pipeline converts raw images to pseudo-colors representing nuclear size: red for larger nuclei and blue for smaller nuclei (**Fig. 1d**). As anticipated, the plot of Paclitaxel-treated cells shifts significantly towards red, indicating an increase in PGCCs, while the untreated cell population predominantly remains blue. Based on the size threshold established in our prior work, we quantify the numbers of PGCCs and non-PGCCs for each image.^38^ This high-throughput screening tool can process up to 10,000 cells per condition within one second, enabling detailed monitoring of cell development and the identification of compounds affecting PGCC populations.

Using the innovative single-cell morphological analysis, we characterized the changes in cell composition when treated with a compound library of 2,726 compounds, each having successfully completed Phase I drug safety confirmation for potential rapid translational impact. One day after cell loading, cells were treated for two days and then stained and imaged to quantify non-PGCCs and PGCCs (**Fig. 2a**). The counts of PGCCs and non-PGCCs were normalized to the numbers in 8 control wells on the same 96-well plate. Among 2,726 compounds, 29 compounds were excluded due to their fluorescent colors that interfere with image processing, and 461 inhibits the total cell number at least by half. However, among those 461 compounds, 236 compounds (51.2%) boosted the number of PGCCs at least by two times. We further examined commonly used chemotherapeutics. We found that Taxanes (Docetaxel and Paclitaxel (Taxol)), Gemcitabine, Carboplatin, Vinorelbine significantly inhibited non-PGCCs but boosted more treatment-resistant PGCCs after treatment. This partially explain why we see an overall tumor shrinkage after treatment, but remaining cancer cells develop therapeutic resistance and relapse in clinics. While Cyclophosphamide monohydrate, Capecitabine, and Fluorouracil do not induce PGCCs, they are not effective in killing cells. The observation clearly highlights the challenges of current chemotherapies in treating TNBC. Given the complicated *in vivo* environment and challenges of effective drug delivery into the core of tumors, the situation will be much worse in patients. As such, our high-throughput screening capability is essential in identifying new compounds that inhibit PGCCs.

**Figure 2.**
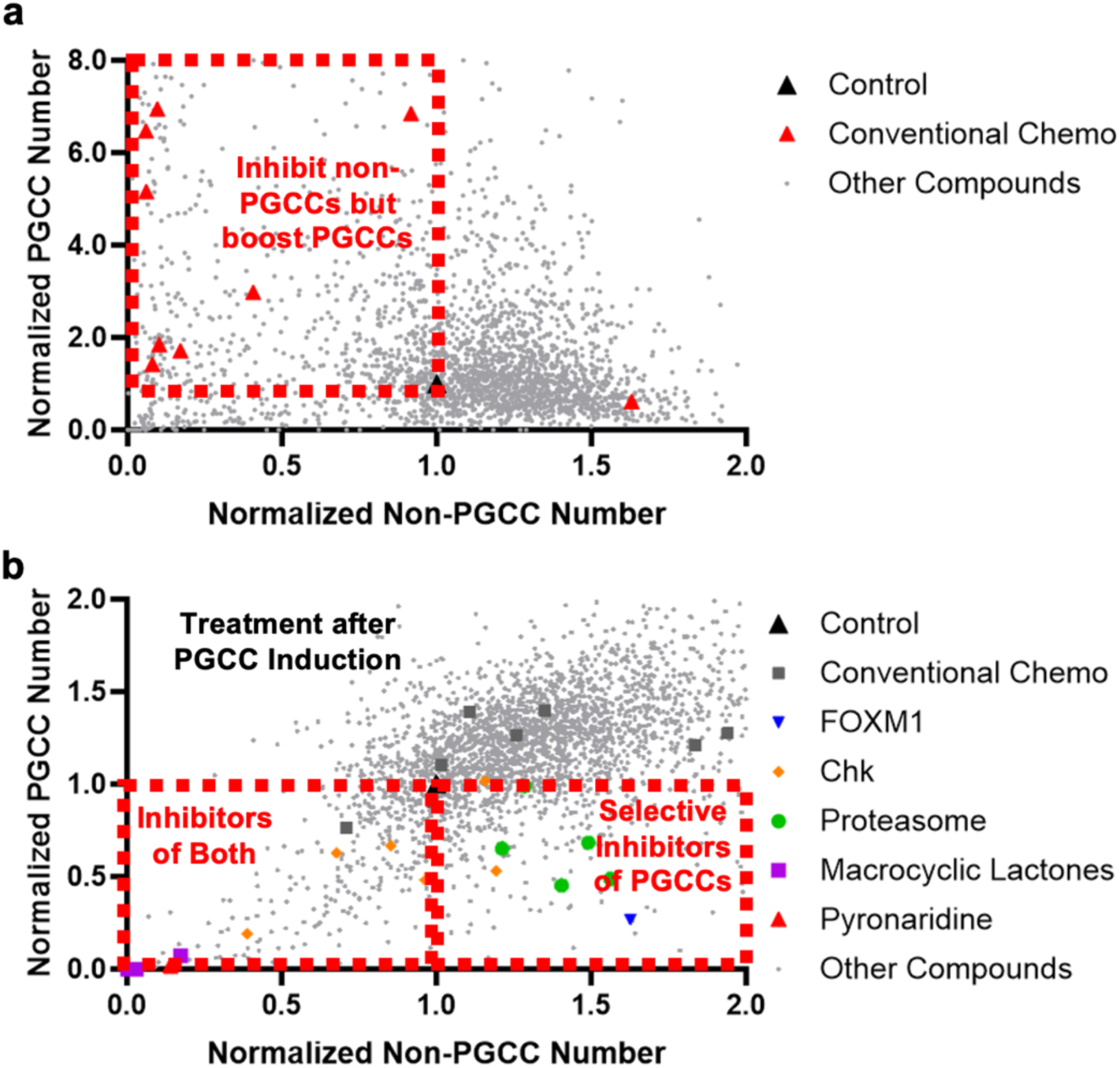
Screening of compounds using MDA-MB-231 breast cancer cells. The X-axis represents the number of non-PGCCs after treatment, and the Y-axis represents the number of PGCCs. Each dot represents the effect of a compound. (a) Direct treatment with screening compounds for 2 days. (b) Pretreatment with 1 µM Paclitaxel for 2 days to induce PGCCs before drug screening.

### Discovering PGCC inhibitors with screening experiments

Given that most TNBC cell lines naturally harbor a minimal PGCC population (<1%), accurately assessing the impact of compounds on PGCCs is challenging. To induce PGCCs, Docetaxel was administered to cells for two days after initial loading.

Subsequently, the cell suspension was aspirated to remove Docetaxel, and the testing compounds were introduced for an additional two days. Cells were then stained and imaged to quantify both PGCCs and non-PGCCs (**Fig. 1a**). As illustrated in **Fig. 2b**, drug-resistant PGCCs proved largely impervious to most compounds. Conventional chemo-therapeutic drugs are also ineffective in killing treatment-resistant PGCCs. Among 2,697 compounds, 169 inhibited PGCCs by at least twofold, 45 inhibited them by at least tenfold, and 63 inhibited both PGCCs and non-PGCCs by at least twofold (**Fig. 2b**).

Among the potent drugs against PGCCs, we observed the efficacy of proteasome inhibitors (e.g., Bortezomib, Oprozomib, Carfilzomib, and Celastrol), CHK inhibitors (e.g., AZD7762, PF-477736), and FOXM1 inhibitor Thiostrepton. FOXM1, a key regulator of the cell cycle, is dysregulated in PGCCs, making them particularly susceptible to FOXM1 inhibition.^38, 61, 62^ Proteasome inhibitors induce cancer cell death through multiple mechanisms, including the accumulation of pro-apoptotic proteins and cell cycle arrest, as well as the buildup of misfolded proteins that heighten cellular stress and sensitivity to other therapies.^63–65^ CHK inhibitors, by targeting CHK1 and CHK2, disrupt DNA damage repair and cell cycle control, preventing cancer cells from recovering from therapy-induced damage and enhancing the efficacy of existing treatments.^66, 67^ While these compounds have been studied, they are not yet in clinical use for treating breast cancer. Their selective activity against PGCCs highlights their potential as therapeutic options for patients with treatment-resistant breast cancer characterized by a significant presence of PGCCs.

In addition to well-studied targets, the large-scale screening revealed promising new compounds for targeting PGCCs (**Fig. 2b**). Notably, macrocyclic lactones such as Ivermectin, Doramectin, and Moxidectin—known for their antiparasitic effects—function by binding to glutamate-gated chloride channels in parasitic nerve and muscle cells.^68–71^ This binding elevates chloride ion permeability, leading to hyperpolarization, paralysis, and death of the parasites. These compounds also interact with other ion channels, disrupting neurotransmission specifically in parasites while leaving host cells largely unaffected due to structural differences in ion channels. Recent studies have demonstrated that Doramectin inhibits glioblastoma cell survival through modulation of autophagy; however, its effects on breast cancer cells have yet to be explored.^72^ Furthermore, Pyronaridine, an antimalarial drug used in combination therapies for Plasmodium falciparum and Plasmodium vivax infections, was also found to effectively eliminate PGCCs.^73, 74^ Pyronaridine’s effect is visually indicated by a blue shift in pseudo color compared to the control (**Fig. 3a**). Pyronaridine disrupts hemozoin formation, leading to toxic heme accumulation, intercalates into DNA to inhibit nucleic acid synthesis, and induces oxidative stress through ROS generation. This multifaceted action damages critical cellular components, killing the parasite. When used with artesunate, Pyronaridine improves treatment efficacy and overcomes resistance, enhancing parasite clearance and therapeutic outcomes. Beyond its antimalarial properties, Doramectin’s antiviral activity against COVID-19 and Ebola viruses has garnered significant attention.^74–76^ Although its potential impact on breast cancer has been noted,^77, 78^ there has been no prior investigation into its ability to overcome therapeutic resistance or specifically target PGCCs. Overall, although the mechanism of PGCC inhibition by these compounds remains unclear, they present intriguing possibilities for future investigation.

**Figure 3.**
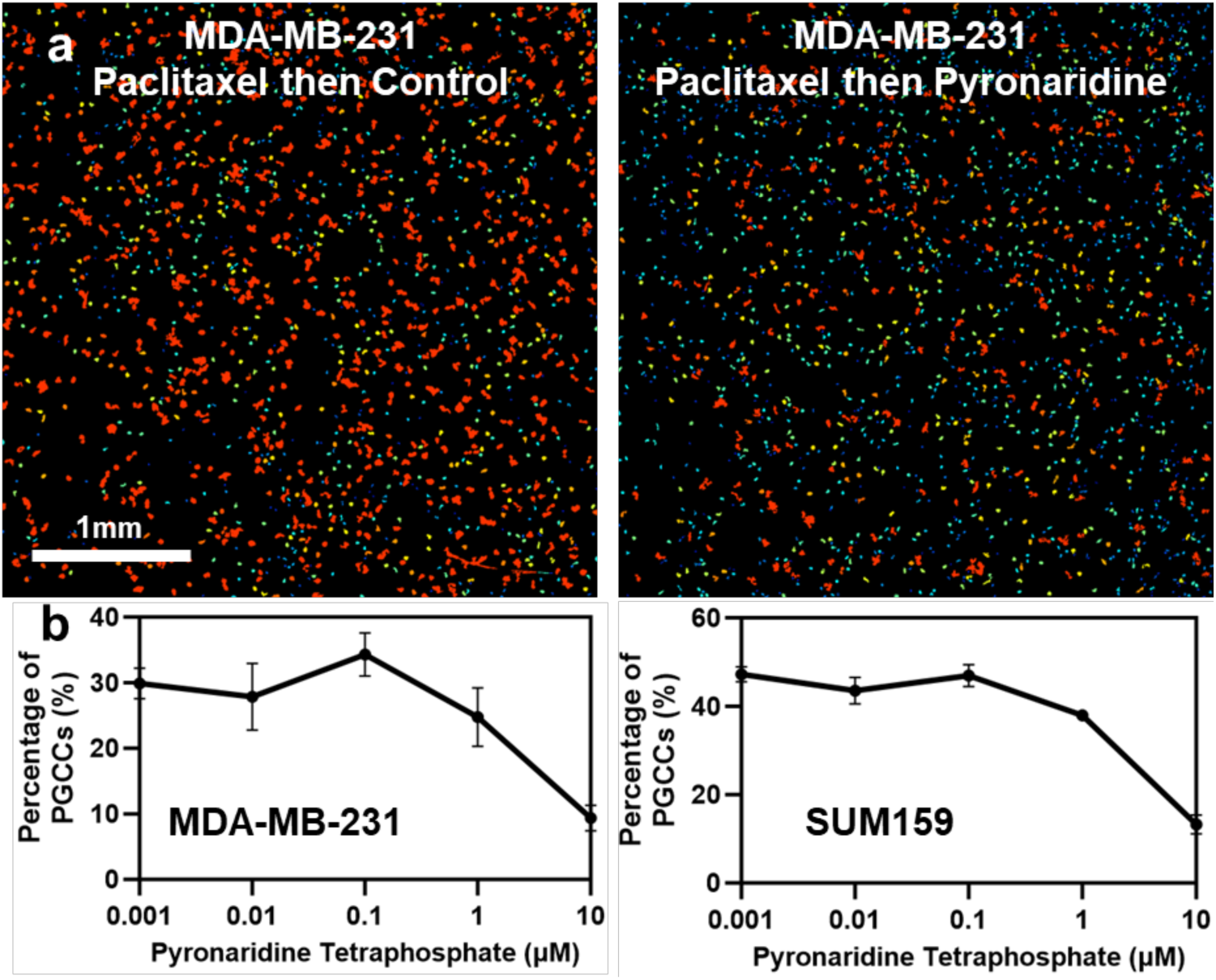
Pyronaridine selectively inhibits PGCCs. (a) Pseudo-color plots indicate that Pyronaridine selectively targets PGCCs, with red representing larger nuclei and blue indicating smaller nuclei. (b) Pyronaridine treatment effects on two TNBC cell lines (MDA-MB-231 and SUM159). The X-axis denotes the compound concentration, while the Y-axis shows the percentage of PGCCs among total cells. Error bars represent the standard deviation; *n* = 4.

In addition to single-dose treatments, we tested five concentrations of selected compounds to validate our screening results in MDA-MB-231 cells (**Fig. 3b**). To further confirm these findings, we evaluated the compounds in a second TNBC line, SUM159. Notably, Pyronaridine selectively targeted PGCCs in both cell types (**Fig. 3b**). These results highlight our distinct capability to differentiate compounds based on their selective effects on PGCCs versus non-PGCCs, enabling precise identification and validation of effective PGCC inhibitors.

### Identification and validation of AXL as a key mediator for the anti-PGCC effects of Pyronaridine

To investigate the potential mechanisms underlying Pyronaridine-induced inhibition of PGCCs in MDA-MB-231 cells, we performed RNA-seq on Pyronaridine-treated PGCCs and compared their gene expression profiles to those of untreated cells. We applied GSEA to identify signaling pathways perturbed by the treatment, focusing on gene sets associated with various perturbations. We identified 283 statistically significantly depleted gene sets (normalized enrichment score [NES] < 0, *q*-value < 0.05) in Pyronaridine-treated cells compared to control cells. In other words, these gene sets were enriched for genes downregulated by Pyronaridine treatment. An association network analysis of these gene sets revealed a close involvement in cell cycle regulation and cancer cell proliferation (**Fig. 4a and b**). Among these gene sets, we observed significant depletion in the KOBAYASHI_EGFR_SIGNALING_24HR_DN, which contains genes downregulated by EGFR inhibition (NES = −1.74, *q* = 0.007) (**Fig. 4a and c**).^79^ This gene set overlapped with several others related to cell cycle states, RB1 targets, and breast cancer grades (**Fig. 4a**). These findings indicate that Pyronaridine may deregulate EGFR signaling pathway to inhibit PGCC proliferation in TNBC, echoing results from a previous report in non-small cell lung cancer.^80^

**Figure 4.**
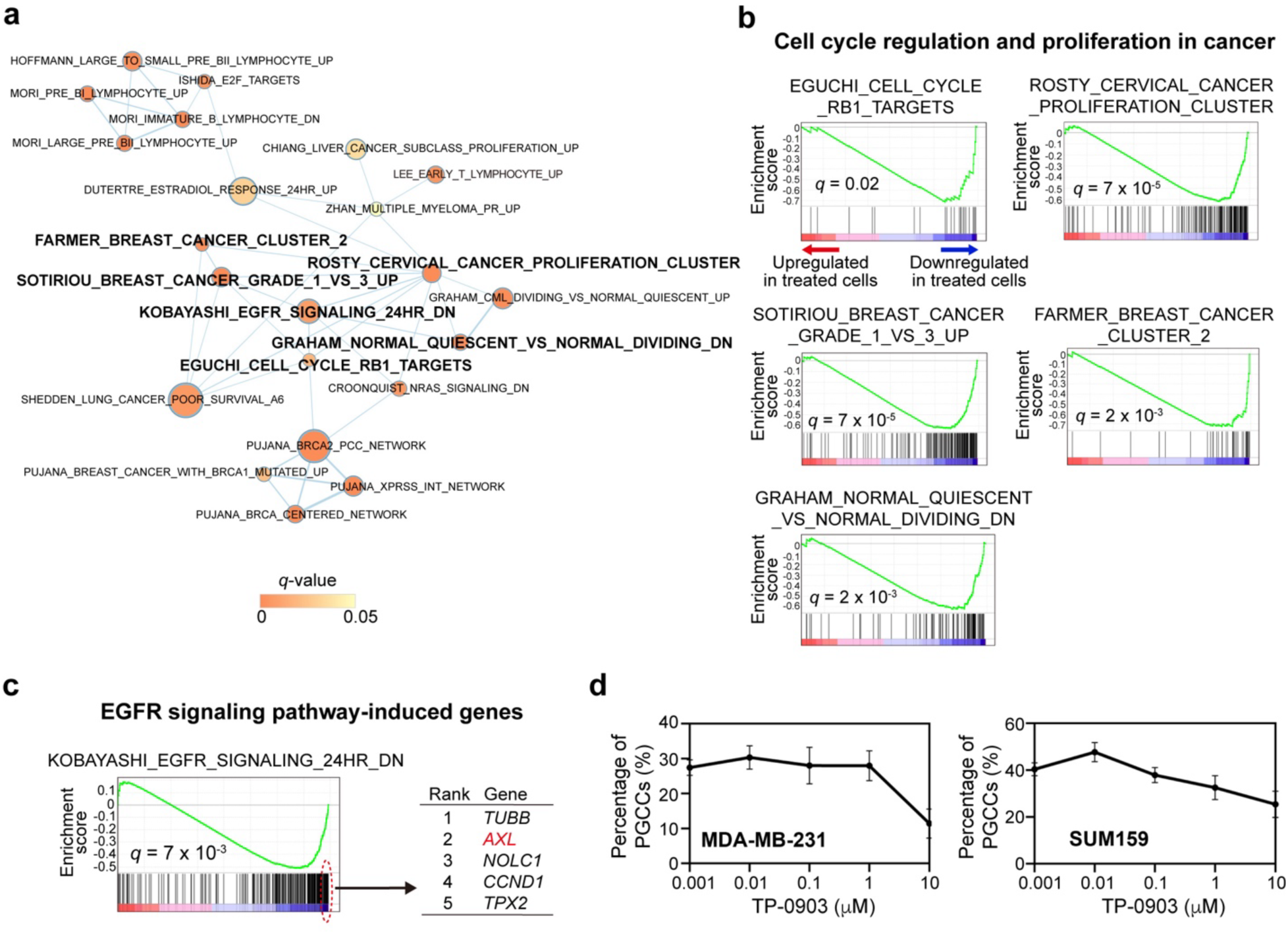
RNA-Seq analysis of Pyronaridine treatment and validation of TP-0903, an AXL inhibitor, in inhibiting PGCCs. (a) GSEA of Pyronaridine-treated cells compared to untreated cells was performed using curated CGP gene sets of MSigDB. A gene set association network was constructed among the significantly depleted gene sets (enriched in Pyronaridine-downregulated genes with NES < 0 and *q*-value < 0.05) and visualized by EnrichmentMap. Each node represents each gene set (node size: gene set size; node color: GSEA *q*-value; edge width: degree of gene overlap between two gene sets [combined coefficient > 0.375]). Gene sets highlighted in bold are further shown in the following panels. (b, c) Significantly depleted gene sets associated with cell cycle regulation and proliferation (b) and EGFR signaling pathway (c) in Pyronaridine-treated cells compared to untreated cells. The 5 top-ranked leading-edge genes in EGFR signaling pathway gene set are shown (c, right panel). (d) Effects of TP-0903 on two TNBC cell lines (MDA-MB-231 and SUM159). The X-axis denotes compound concentration, and the Y-axis shows the percentage of PGCCs among total cells. Error bars represent the standard deviation; *n* = 4.

We further explored key players in the EGFR signaling pathway-mediated genes for their potential as therapeutic targets of PGCCs in TNBC. The 5 top-ranked leading-edge genes from GSEA included *TUBB*, *AXL*, *NOLC1*, *CCND1*, and *TPX2* (**Fig. 4c**), all of which were significantly downregulated in Pyronaridine-treated cells. Among these, AXL emerged as a particularly promising target for further investigation. The AXL pathway, driven by the AXL receptor tyrosine kinase, orchestrates cell survival, proliferation, migration, and invasion.^81–83^ Activation by its ligand, Gas6, triggers a signaling cascade involving PI3K, AKT, and MAPK, which enhances cell survival, inhibits apoptosis, promotes epithelial-to-mesenchymal transition (EMT), and facilitates cancer metastasis.^84^ AXL also plays a role in immune evasion and therapy resistance, with its dysregulation often correlating with aggressive cancer phenotypes and poor prognosis, making it a prime target for therapeutic intervention.^85^ In light of our RNA-Seq data and existing literature on AXL’s role in therapy resistance, we tested TP-0903, a novel, orally bioavailable AXL inhibitor currently in a first-in-human clinical trial for advanced solid tumors.^86, 87^ As an ATP-competitive inhibitor, it features an adenine-mimicking heterocyclic structure and specifically binds to the active form of AXL. Our findings demonstrate that TP-0903 effectively targets PGCCs in both MDA-MB-231 and SUM159 cells (**Fig. 4d**). This preliminary study aligns with RNA-Seq analysis and suggests that Pyronaridine’s mechanism in targeting PGCCs may involve the AXL pathway.

### Machine learning-based prediction of anti-PGCC effects using high-throughput screening data

The impracticality of empirically screening all existing compounds and the absence of predictive models are major obstacles hindering the identification of promising anti-PGCC compounds for experimental validation. To address this challenge, we assessed the potential of our high-throughput morphological assay of 2,726 compounds to effectively inform predictive machine learning models. Specifically, we comprehensively tested seven state-of-the-art machine learning methods to predict anti-PGCC efficacy in MDA-MB-231 cells. As described in Methods, these regression models were trained to predict changes in PGCC counts based on quantitative representations of either chemical structures (fingerprints) or compound descriptions (text converted to embeddings) (**Fig. 5a**). A total of 2,430 compounds in the screening library with both features available were used in the model. We adopted 10 rounds of 10-fold cross-validations to train and test each model. In each iteration of cross-validation, a model was trained using 90% of the 2,430 compounds and tested on the remaining 10%, which were not seen by the model during training. Overall, 31 out of 63 (49.2%) models achieved a median Pearson correlation coefficient *ρ* above 0.2 across 10 rounds of cross-validations (**Fig. 5b**).

**Figure 5.**
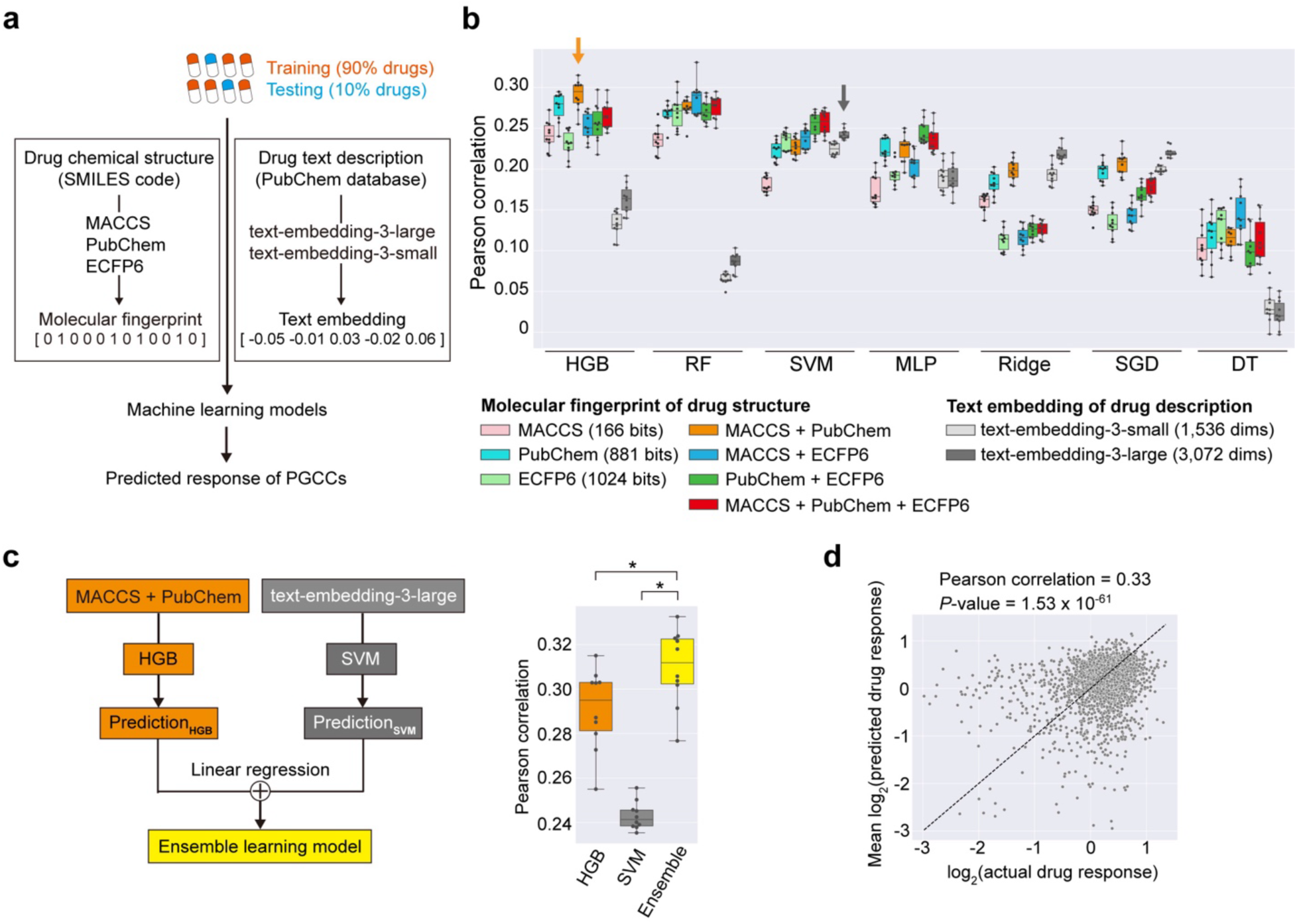
Machine learning models for predicting anti-PGCC effects of compounds. (a) Schematic of machine learning models used to predict drug responses of PGCCs in MDA-MB-231 cells based on drug chemical structures and descriptions, represented by fingerprint and embedding vectors, respectively. A total of 2,430 compounds in the screening library with both features available were used for training and testing the model (2,187 for training and 243 for testing). (b) Predictive performance comparisons among seven state-of-the-art machine learning models trained using either single or multiple molecular fingerprints, or a text embedding. The performance is measured by the Pearson correlation coefficient between actual and predicted drug response values. Ten rounds of 10-fold cross-validation were performed to train and test each model. Each dot in the box plots represents the average of 10 correlation coefficients obtained from 10-fold cross-validations in each round. Orange and grey arrows indicate the best-performing models for molecular fingerprints and text embeddings, respectively, selected for the ensemble learning model. Abbreviations: dims, dimensions; HGB, histogram-based gradient boosting; RF, random forest; SVM, support vector machine; MLP, multi-layer perceptron; Ridge, linear regression with L2 regularization; SGD, stochastic gradient descent linear regression; DT, decision tree. (c) Left panel: schematic of an ensemble learning model trained by integrating the best-performing models for drug structures and descriptions. Right panel: predictive performance comparison of the ensemble learning model versus individual models (*: one-tailed paired t-test *P* < 1×10^-6^). (d) Predictive performance of the ensemble model across all 2,430 drugs. For each drug, the average predicted response across the 10 rounds is shown in the plot. Only drugs with log2-transformed actual and predicted response values greater than −3 are shown in the plot.

For molecular fingerprints, HGB with a combination of MACCS and PubChem was the best model (median *ρ*, 0.29; **Fig. 5b**). Models that used combinations of multiple molecular fingerprints as features tended to achieve better performance compared to those using single molecular fingerprints. For example, HGB with MACCS and PubChem, RF with MACCS and ECFP6, and SVM with all three molecular fingerprints outperformed their single-fingerprint counterparts (**Fig. 5b**). For description-based embeddings, models with longer embeddings (3,072 dimensions) generally outperformed those with 1,536 dimensions (**Fig. 5b**), suggesting that longer embeddings capture additional pharmacological information. Notably, SVM with 3,072-dimensional embeddings was the best-performing model (median *ρ* = 0.24; **Fig. 5b**). Overall, performance of these models was comparable to the best results from a community challenge for predicting drug sensitivities and recent studies predicting genetic dependencies in pan-cancer cell lines,^88–90^ demonstrating the capability of our screening library to support accurate predictive modeling.

### Enhancing predictive performance by integrating compound structures and descriptions using an ensemble learning approach

Since compound structures and descriptions provide distinct yet potentially complementary information, combining these features may improve the performance of predictive models. To explore this, we developed an ensemble learning method by integrating the best-performing models for drug structures and descriptions, respectively (*i.e.*, HGB on MACCS and PubChem, and SVM on the longer embedding). The ensemble model utilized linear regression to generate the final prediction based on the outputs of these two models. Notably, this approach significantly improved performance (median *ρ* = 0.31) compared to the individual models (one-tailed paired *t*-test, both *P* < 1×10^-6^) (**Fig. 5c**). Across all 2,430 drugs, the ensemble model achieved a *ρ* of 0.33 between real and predicted drug responses (*P* = 1.53 x 10^-61^) (**Fig. 5d**).

In the ensemble model, the regression coefficients for the HGB and SVM models were 1.2 and 0.6, respectively, both statistically significant (*P* < 1×10^-3^). These results suggest that both models contributed meaningful and independent information to the ensemble model. The HGB model had a greater impact on the final prediction, while the SVM model predictions provided a complementary effect. Taken together, our findings demonstrate that integrating these two distinct features allows the model to capture meaningful and complementary patterns related to anti-PGCC effects, leading to enhanced predictive performance.

### Expanded virtual screening by the ensemble prediction model and validation using a patient-derived model

We expanded our virtual screening to a broader range of compounds to identify potential anti-PGCC agents in breast cancer. We compiled a large library of compounds based on the Profiling Relative Inhibition Simultaneously in Mixtures (PRISM) project, which is one of the largest drug sensitivity screens, covering 6,575 oncology or non-oncology drugs (as of 24Q2).^91^ Of these 6,575 drugs, 3,093 drugs were not part of our original screening library but had both drug structure and description information. We applied our ensemble model to predict anti-PGCC effects for these 3,093 drugs in MDA-MB-231 cells. The predicted drug rankings, based on their viability-inhibitory effects in PGCCs, are shown in **Fig. 6a**. Among the top-ranked candidates, we prioritized those with novelty, strong pharmacological profiles, and translational potential for experimental validation. Notably, two compounds—Lestaurtinib and UCN-01—demonstrated effective inhibition of PGCCs in both MDA-MB-231 and SUM159 cell lines, validating the model’s predictions (**Fig. 6b**).

**Figure 6.**
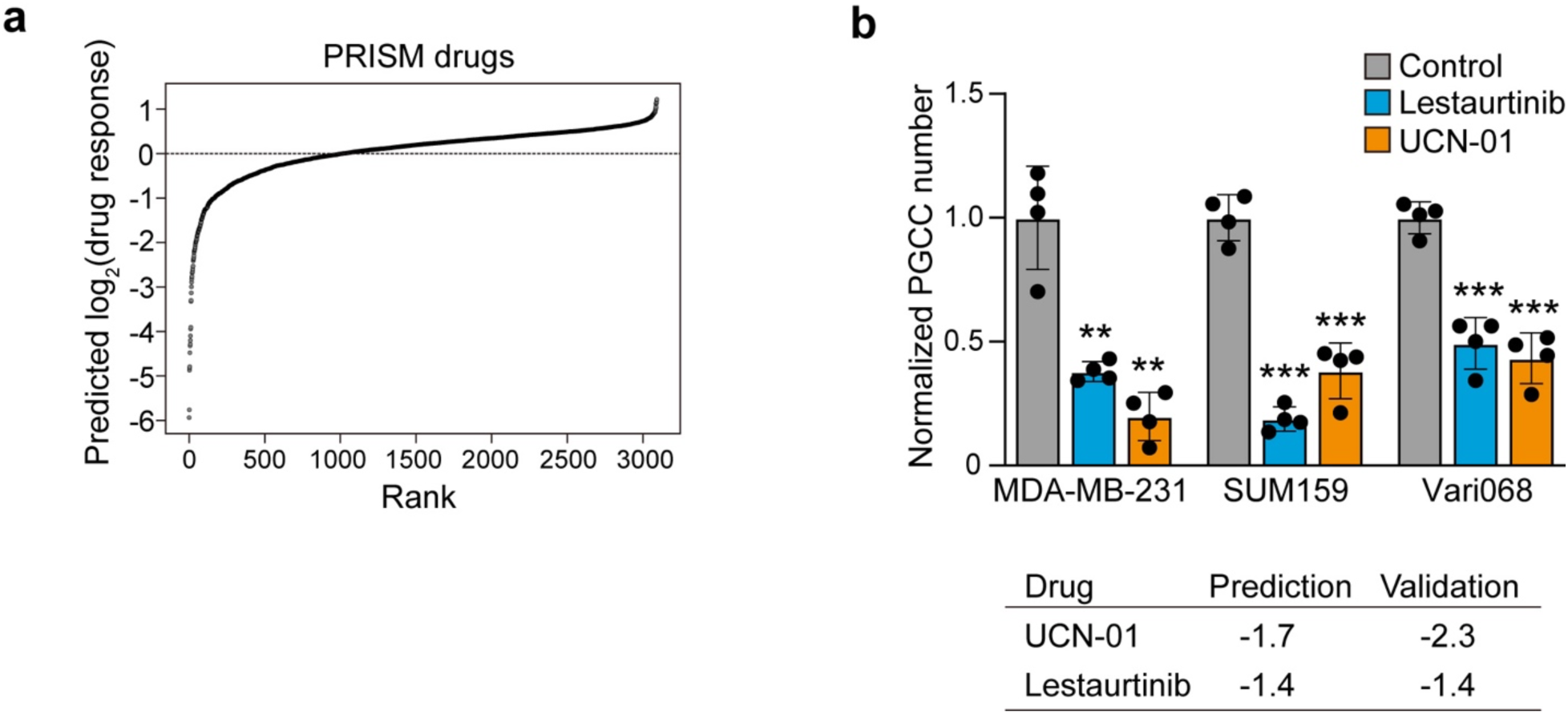
Compound candidates (Lestaurtinib and UCN-01) predicted by ensemble learning model effectively inhibit PGCCs. (a) Rank plot of predicted log₂-transformed drug response values in PGCCs of MDA-MB-231 cells, using our ensemble learning model based on drugs included in the PRISM project. (b) Validation of the two compounds on two TNBC cell lines (MDA-MB-231 and SUM159) and low-passage patient-derived Vari068 cells. The number of PGCCs were normalized to the control, as per the training data format. Error bars represent the standard deviation; *n* = 4. ** indicates *P* < 0.01, and *** indicates *p* < 0.001. The lower panel shows the predicted and validated drug responses for UCN-01 and Lestaurtinib.

To further ensure the clinical relevance of these findings, we validated the two compounds in a low-passage, TNBC patient-derived cell line, Vari068, which naturally harbors a high population of PGCCs. Remarkably, we confirmed a significant reduction in PGCCs within this patient-derived model. Although machine learning models do not always provide direct mechanistic explanations, a literature review suggests plausible mechanisms. Lestaurtinib, a multi-targeted tyrosine kinase inhibitor, interferes with stress signaling pathways involving JAK2, which PGCCs depend on for survival.^92–95^ UCN-01, a Chk1 inhibitor, targets crucial cell cycle checkpoints, undermining PGCCs’ ability to manage DNA damage and genomic instability.^96, 97^ By disrupting these survival pathways, both drugs induce PGCC vulnerability, leading to selective cell death. The successful validation of these model-predicted compounds demonstrates the significant potential of machine learning-based virtual screening to accelerate the discovery of novel anti-cancer therapies, particularly for targeting therapy-resistant PGCCs.

## Conclusions

Therapy resistance in breast cancer is increasingly linked to the presence of PGCCs, which arise through whole-genome doubling and exhibit heightened resistance to conventional treatments. To address the challenge of identifying effective PGCC inhibitors in a high-throughput manner, we developed a single-cell morphological analysis workflow that rapidly distinguishes compounds targeting non-PGCCs, PGCCs, or both. Through screening a library of 2,726 FDA Phase 1-approved drugs, we identified several promising anti-PGCC candidates, including inhibitors of the proteasome, FOXM1, CHK, and macrocyclic lactones. RNA-Seq analysis of Pyronaridine-treated cells further suggested that AXL inhibition could be a viable strategy for targeting PGCCs. To scale up the discovery of potential PGCC inhibitors, we developed an ensemble learning model that predicts anti-PGCC efficacy by integrating two machine learning models based on chemical fingerprints and compound descriptions. This model successfully predicted effective compounds from the PRISM library, which includes over 6,000 drugs. Two of the top-ranked predictions were experimentally validated as potent PGCC inhibitors. These findings underscore the potential of machine learning-driven virtual screening to accelerate the discovery of novel therapies aimed at overcoming therapy resistance in PGCCs.

## Author Contributions

Drug screening and cell biology experiments were performed by Jinxiong Cheng, Hsiao-Chun Chen, and Yu-Chih Chen. Software for single-cell morphological analysis was developed by Yushu Ma. RNA-Seq experiment was performed by Yu-Chih Chen. Sequencing read alignment and data analysis were performed by Chien-Hung Shih and Yu-Chih Chen. *In silico* prediction of PGCC inhibitors was performed by Chien-Hung Shih, Li-Ju Wang, Yanhao Tan, and Yu-Chiao Chiu. Yu-Chih Chen and Yu-Chiao Chiu supervised the study. Chien-Hung Shih, Yu-Chiao Chiu, and Yu-Chih Chen wrote the manuscript. All authors discussed the results, commented on the manuscript, and approved the final manuscript.

## Conflicts of Interest

The authors declare no competing interests.

## Acknowledgements

This study was generously funded by start-up support from the UPMC Hillman Cancer Center awarded to Yu-Chih Chen and Yu-Chiao Chiu (supported by the National Institutes of Health [NIH] through Grant Numbers P30CA047904 and P50CA272218), the Women’s Cancer Research Center (WCRC) at Magee Women’s Research Institute to Yu-Chih Chen, the Pitt CTSI Pilot project to Yu-Chih Chen (NIH Grant Number UL1TR001857), Pittsburgh Liver Research Center (NIH P30DK120531) to Yu-Chiao Chiu, the NIH National Institute of General Medical Sciences (R35GM150509 to Yu-Chih Chen and R35GM154967 to Yu-Chiao Chiu), the NIH National Cancer Institute to Yu-Chiao Chiu (R00CA248944), the NIH Office of the Director to Yu-Chiao Chiu (3R00CA248944-04S1 and R03DE033361), and Leukemia Research Foundation to Yu-Chiao Chiu, as well as the UPMC Competitive Medical Research Fund (CMRF) awarded to Yu-Chih Chen. This research was supported in part by the University of Pittsburgh Center for Research Computing (NIH S10OD028483), through the resources provided. We also thank Drs. Gary Luker and Max Wicha at the University of Michigan for kindly providing the cell lines used in this study.

## Declaration of Generative AI in Scientific Writing

The authors utilized ChatGPT (versions, 4o and 3.5) to enhance the readability and language of this work. Following its use, the authors thoroughly reviewed and edited the content as necessary and take full responsibility for the content of the publication.

## Notes

### Competing Interest Statement

The authors have declared no competing interest.

## References

(1) Comai, L. The advantages and disadvantages of being polyploid. Nat Rev Genet 2005, 6 (11), 836–846. DOI: 10.1038/nrg1711.

(2) Ogden, A.; Rida, P. C.; Knudsen, B. S.; Kucuk, O.; Aneja, R. Docetaxel-induced polyploidization may underlie chemoresistance and disease relapse. Cancer Lett 2015, 367 (2), 89–92. DOI: 10.1016/j.canlet.2015.06.025.

(3) Bharadwaj, D.; Mandal, M. Senescence in polyploid giant cancer cells: A road that leads to chemoresistance. Cytokine Growth Factor Rev 2020, 52, 68–75. DOI: 10.1016/j.cytogfr.2019.11.002.

(4) Garrido Castillo, L. N.; Anract, J.; Delongchamps, N. B.; Huillard, O.; BenMohamed, F.; Decina, A.; Lebret, T.; Dachez, R.; Paterlini-Brechot, P. Polyploid Giant Cancer Cells Are Frequently Found in the Urine of Prostate Cancer Patients. Cancers (Basel) 2023, 15 (13). DOI: 10.3390/cancers15133366.

(5) Zhou, X.; Zhou, M.; Zheng, M.; Tian, S.; Yang, X.; Ning, Y.; Li, Y.; Zhang, S. Polyploid giant cancer cells and cancer progression. Front Cell Dev Biol 2022, 10, 1017588. DOI: 10.3389/fcell.2022.1017588.

(6) Richards, J. S.; Candelaria, N. R.; Lanz, R. B. Polyploid giant cancer cells and ovarian cancer: new insights into mitotic regulators and polyploidydagger. Biol Reprod 2021, 105 (2), 305–316. DOI: 10.1093/biolre/ioab102.

(7) Bowers, R. R.; Andrade, M. F.; Jones, C. M.; White-Gilbertson, S.; Voelkel-Johnson, C.; Delaney, J. R. Autophagy modulating therapeutics inhibit ovarian cancer colony generation by polyploid giant cancer cells (PGCCs). BMC Cancer 2022, 22 (1), 410. DOI: 10.1186/s12885-022-09503-6.

(8) Bai, S.; Taylor, S. E.; Jamalruddin, M. A.; McGonigal, S.; Grimley, E.; Yang, D.; Bernstein, K. A.; Buckanovich, R. J. Targeting Therapeutic Resistance and Multinucleate Giant Cells in CCNE1-Amplified HR-Proficient Ovarian Cancer. Mol Cancer Ther 2022, 21 (9), 1473–1484. DOI: 10.1158/1535-7163.MCT-21-0873.

(9) Mannan, R.; Wang, X.; Bawa, P. S.; Spratt, D. E.; Wilson, A.; Jentzen, J.; Chinnaiyan, A. M.; Reichert, Z. R.; Mehra, R. Polypoidal giant cancer cells in metastatic castration-resistant prostate cancer: observations from the Michigan Legacy Tissue Program. Med Oncol 2020, 37 (3), 16. DOI: 10.1007/s12032-020-1341-6.

(10) Qu, Y.; Zhang, L.; Rong, Z.; He, T.; Zhang, S. Number of glioma polyploid giant cancer cells (PGCCs) associated with vasculogenic mimicry formation and tumor grade in human glioma. J Exp Clin Cancer Res 2013, 32, 75. DOI: 10.1186/1756-9966-32-75.

(11) Pienta, K. J.; Hammarlund, E. U.; Axelrod, R.; Brown, J. S.; Amend, S. R. Poly-aneuploid cancer cells promote evolvability, generating lethal cancer. Evol Appl 2020, 13 (7), 1626–1634. DOI: 10.1111/eva.12929.

(12) Xuan, B.; Ghosh, D.; Cheney, E. M.; Clifton, E. M.; Dawson, M. R. Dysregulation in Actin Cytoskeletal Organization Drives Increased Stiffness and Migratory Persistence in Polyploidal Giant Cancer Cells. Sci Rep 2018, 8 (1), 11935. DOI: 10.1038/s41598-018-29817-5.

(13) Mosieniak, G.; Sliwinska, M. A.; Alster, O.; Strzeszewska, A.; Sunderland, P.; Piechota, M.; Was, H.; Sikora, E. Polyploidy Formation in Doxorubicin-Treated Cancer Cells Can Favor Escape from Senescence. Neoplasia 2015, 17 (12), 882–893. DOI: 10.1016/j.neo.2015.11.008.

(14) Quinton, R. J.; DiDomizio, A.; Vittoria, M. A.; Kotynkova, K.; Ticas, C. J.; Patel, S.; Koga, Y.; Vakhshoorzadeh, J.; Hermance, N.; Kuroda, T. S.;, et al. Whole-genome doubling confers unique genetic vulnerabilities on tumour cells. Nature 2021, 590 (7846), 492–497. DOI: 10.1038/s41586-020-03133-3.

(15) Saini, G.; Joshi, S.; Garlapati, C.; Li, H.; Kong, J.; Krishnamurthy, J.; Reid, M. D.; Aneja, R. Polyploid giant cancer cell characterization: New frontiers in predicting response to chemotherapy in breast cancer. Semin Cancer Biol 2022, 81, 220–231. DOI: 10.1016/j.semcancer.2021.03.017.

(16) Stephens, P. J.; Greenman, C. D.; Fu, B.; Yang, F.; Bignell, G. R.; Mudie, L. J.; Pleasance, E. D.; Lau, K. W.; Beare, D.; Stebbings, L. A.;, et al. Massive genomic rearrangement acquired in a single catastrophic event during cancer development. Cell 2011, 144 (1), 27–40. DOI: 10.1016/j.cell.2010.11.055.

(17) Mittal, K.; Donthamsetty, S.; Kaur, R.; Yang, C.; Gupta, M. V.; Reid, M. D.; Choi, D. H.; Rida, P. C. G.; Aneja, R. Multinucleated polyploidy drives resistance to Docetaxel chemotherapy in prostate cancer. Br J Cancer 2017, 116 (9), 1186–1194. DOI: 10.1038/bjc.2017.78.

(18) Fei, F.; Zhang, D.; Yang, Z.; Wang, S.; Wang, X.; Wu, Z.; Wu, Q.; Zhang, S. The number of polyploid giant cancer cells and epithelial-mesenchymal transition-related proteins are associated with invasion and metastasis in human breast cancer. J Exp Clin Cancer Res 2015, 34, 158. DOI: 10.1186/s13046-015-0277-8.

(19) Wang, X.; Zheng, M.; Fei, F.; Li, C.; Du, J.; Liu, K.; Li, Y.; Zhang, S. EMT-related protein expression in polyploid giant cancer cells and their daughter cells with different passages after triptolide treatment. Med Oncol 2019, 36 (9), 82. DOI: 10.1007/s12032-019-1303-z.

(20) Zhang, S.; Mercado-Uribe, I.; Xing, Z.; Sun, B.; Kuang, J.; Liu, J. Generation of cancer stem-like cells through the formation of polyploid giant cancer cells. Oncogene 2014, 33 (1), 116–128. DOI: 10.1038/onc.2013.96.

(21) Gerashchenko, B. I.; Salmina, K.; Eglitis, J.; Huna, A.; Grjunberga, V.; Erenpreisa, J. Disentangling the aneuploidy and senescence paradoxes: a study of triploid breast cancers non-responsive to neoadjuvant therapy. Histochem Cell Biol 2016, 145 (4), 497–508. DOI: 10.1007/s00418-016-1415-x.

(22) Salmina, K.; Jankevics, E.; Huna, A.; Perminov, D.; Radovica, I.; Klymenko, T.; Ivanov, A.; Jascenko, E.; Scherthan, H.; Cragg, M.;, et al. Up-regulation of the embryonic self-renewal network through reversible polyploidy in irradiated p53-mutant tumour cells. Exp Cell Res 2010, 316 (13), 2099–2112. DOI: 10.1016/j.yexcr.2010.04.030.

(23) Zhang, X.; Yao, J.; Li, X.; Niu, N.; Liu, Y.; Hajek, R. A.; Peng, G.; Westin, S.; Sood, A. K.; Liu, J. Targeting polyploid giant cancer cells potentiates a therapeutic response and overcomes resistance to PARP inhibitors in ovarian cancer. Sci Adv 2023, 9 (29), eadf7195. DOI: 10.1126/sciadv.adf7195.

(24) Vicente, J. J.; Khan, K.; Tillinghast, G.; McFaline-Figueroa, J. L.; Sancak, Y.; Stella, N. The microtubule targeting agent ST-401 triggers cell death in interphase and prevents the formation of polyploid giant cancer cells. J Transl Med 2024, 22 (1), 441. DOI: 10.1186/s12967-024-05234-3.

(25) Adibi, R.; Moein, S.; Gheisari, Y. Zoledronic acid targets chemo-resistant polyploid giant cancer cells. Sci Rep 2023, 13 (1), 419. DOI: 10.1038/s41598-022-27090-1.

(26) White-Gilbertson, S.; Lu, P.; Saatci, O.; Sahin, O.; Delaney, J. R.; Ogretmen, B.; Voelkel-Johnson, C. Transcriptome analysis of polyploid giant cancer cells and their progeny reveals a functional role for p21 in polyploidization and depolyploidization. J Biol Chem 2024, 300 (4), 107136. DOI: 10.1016/j.jbc.2024.107136.

(27) White-Gilbertson, S.; Lu, P.; Esobi, I.; Echesabal-Chen, J.; Mulholland, P. J.; Gooz, M.; Ogretmen, B.; Stamatikos, A.; Voelkel-Johnson, C. Polyploid giant cancer cells are dependent on cholesterol for progeny formation through amitotic division. Sci Rep 2022, 12 (1), 8971. DOI: 10.1038/s41598-022-12705-4.

(28) You, B.; Xia, T.; Gu, M.; Zhang, Z.; Zhang, Q.; Shen, J.; Fan, Y.; Yao, H.; Pan, S.; Lu, Y.;, et al. AMPK-mTOR-Mediated Activation of Autophagy Promotes Formation of Dormant Polyploid Giant Cancer Cells. Cancer Res 2022, 82 (5), 846–858. DOI: 10.1158/0008-5472.CAN-21-2342.

(29) Lissa, D.; Senovilla, L.; Rello-Varona, S.; Vitale, I.; Michaud, M.; Pietrocola, F.; Boileve, A.; Obrist, F.; Bordenave, C.; Garcia, P.;, et al. Resveratrol and aspirin eliminate tetraploid cells for anticancer chemoprevention. Proc Natl Acad Sci U S A 2014, 111 (8), 3020–3025. DOI: 10.1073/pnas.1318440111.

(30) White-Gilbertson, S.; Lu, P.; Jones, C. M.; Chiodini, S.; Hurley, D.; Das, A.; Delaney, J. R.; Norris, J. S.; Voelkel-Johnson, C. Tamoxifen is a candidate first-in-class inhibitor of acid ceramidase that reduces amitotic division in polyploid giant cancer cells-Unrecognized players in tumorigenesis. Cancer Med 2020, 9 (9), 3142–3152. DOI: 10.1002/cam4.2960.

(31) Senovilla, L.; Vitale, I.; Martins, I.; Tailler, M.; Pailleret, C.; Michaud, M.; Galluzzi, L.; Adjemian, S.; Kepp, O.; Niso-Santano, M.;, et al. An immunosurveillance mechanism controls cancer cell ploidy. Science 2012, 337 (6102), 1678–1684. DOI: 10.1126/science.1224922.

(32) Boileve, A.; Senovilla, L.; Vitale, I.; Lissa, D.; Martins, I.; Metivier, D.; van den Brink, S.; Clevers, H.; Galluzzi, L.; Castedo, M.;, et al. Immunosurveillance against tetraploidization-induced colon tumorigenesis. Cell Cycle 2013, 12 (3), 473–479. DOI: 10.4161/cc.23369.

(33) Long, F. Microscopy cell nuclei segmentation with enhanced U-Net. BMC Bioinformatics 2020, 21 (1), 8. DOI: 10.1186/s12859-019-3332-1.

(34) Garvey, C. M.; Spiller, E.; Lindsay, D.; Chiang, C. T.; Choi, N. C.; Agus, D. B.; Mallick, P.; Foo, J.; Mumenthaler, S. M. A high-content image-based method for quantitatively studying context-dependent cell population dynamics. Sci Rep 2016, 6, 29752. DOI: 10.1038/srep29752.

(35) Mzurikwao, D.; Khan, M. U.; Samuel, O. W.; Cinatl, J., Jr.; Wass, M.; Michaelis, M.; Marcelli, G.; Ang, C. S. Towards image-based cancer cell lines authentication using deep neural networks. Sci Rep 2020, 10 (1), 19857. DOI: 10.1038/s41598-020-76670-6.

(36) Pachitariu, M.; Stringer, C. Cellpose 2.0: how to train your own model. Nat Methods 2022, 19 (12), 1634–1641. DOI: 10.1038/s41592-022-01663-4.

(37) He, S.; Sillah, M.; Cole, A. R.; Uboveja, A.; Aird, K. M.; Chen, Y. C.; Gong, Y. N. D-MAINS: A Deep-Learning Model for the Label-Free Detection of Mitosis, Apoptosis, Interphase, Necrosis, and Senescence in Cancer Cells. Cells 2024, 13 (12). DOI: 10.3390/cells13121004.

(38) Zhou, M.; Ma, Y.; Chiang, C. C.; Rock, E. C.; Butler, S. C.; Anne, R.; Yatsenko, S.; Gong, Y.; Chen, Y. C. Single-cell morphological and transcriptome analysis unveil inhibitors of polyploid giant breast cancer cells in vitro. Commun Biol 2023, 6 (1), 1301. DOI: 10.1038/s42003-023-05674-5.

(39) Chiu, Y. C.; Chen, H. H.; Zhang, T.; Zhang, S.; Gorthi, A.; Wang, L. J.; Huang, Y.; Chen, Y. Predicting drug response of tumors from integrated genomic profiles by deep neural networks. BMC Med Genomics 2019, 12 (Suppl 1), 18. DOI: 10.1186/s12920-018-0460-9 From NLM Medline.

(40) Chawla, S.; Rockstroh, A.; Lehman, M.; Ratther, E.; Jain, A.; Anand, A.; Gupta, A.; Bhattacharya, N.; Poonia, S.; Rai, P.;, et al. Gene expression based inference of cancer drug sensitivity. Nat Commun 2022, 13 (1), 5680. DOI: 10.1038/s41467-022-33291-z.

(41) Kuenzi, B. M.; Park, J.; Fong, S. H.; Sanchez, K. S.; Lee, J.; Kreisberg, J. F.; Ma, J.; Ideker, T. Predicting Drug Response and Synergy Using a Deep Learning Model of Human Cancer Cells. Cancer Cell 2020, 38 (5), 672–684 e676. DOI: 10.1016/j.ccell.2020.09.014.

(42) Park, S.; Silva, E.; Singhal, A.; Kelly, M. R.; Licon, K.; Panagiotou, I.; Fogg, C.; Fong, S.; Lee, J. J. Y.; Zhao, X.;, et al. A deep learning model of tumor cell architecture elucidates response and resistance to CDK4/6 inhibitors. Nat Cancer 2024, 5 (7), 996–1009. DOI: 10.1038/s43018-024-00740-1.

(43) Gerdes, H.; Casado, P.; Dokal, A.; Hijazi, M.; Akhtar, N.; Osuntola, R.; Rajeeve, V.; Fitzgibbon, J.; Travers, J.; Britton, D.;, et al. Drug ranking using machine learning systematically predicts the efficacy of anti-cancer drugs. Nat Commun 2021, 12 (1), 1850. DOI: 10.1038/s41467-021-22170-8.

(44) Zhao, H.; Zhang, X.; Zhao, Q.; Li, Y.; Wang, J. MSDRP: a deep learning model based on multisource data for predicting drug response. Bioinformatics 2023, 39 (9). DOI: 10.1093/bioinformatics/btad514.

(45) Pham, T. H.; Qiu, Y.; Liu, J.; Zimmer, S.; O’Neill, E.; Xie, L.; Zhang, P. Chemical-induced gene expression ranking and its application to pancreatic cancer drug repurposing. Patterns (N Y) 2022, 3 (4), 100441. DOI: 10.1016/j.patter.2022.100441.

(46) Liu, M.; Liu, Y.; Deng, L.; Wang, D.; He, X.; Zhou, L.; Wicha, M. S.; Bai, F.; Liu, S. Transcriptional profiles of different states of cancer stem cells in triple-negative breast cancer. Mol Cancer 2018, 17 (1), 65. DOI: 10.1186/s12943-018-0809-x.

(47) Aw Yong, K. M.; Ulintz, P. J.; Caceres, S.; Cheng, X.; Bao, L.; Wu, Z.; Jiagge, E. M.; Merajver, S. D. Heterogeneity at the invasion front of triple negative breast cancer cells. Sci Rep 2020, 10 (1), 5781. DOI: 10.1038/s41598-020-62516-8.

(48) Chen, Y. C.; Ingram, P. N.; Fouladdel, S.; McDermott, S. P.; Azizi, E.; Wicha, M. S.; Yoon, E. High-Throughput Single-Cell Derived Sphere Formation for Cancer Stem-Like Cell Identification and Analysis. Sci Rep 2016, 6, 27301. DOI: 10.1038/srep27301.

(49) Al-Kofahi, Y.; Lassoued, W.; Lee, W.; Roysam, B. Improved automatic detection and segmentation of cell nuclei in histopathology images. IEEE Trans Biomed Eng 2010, 57 (4), 841–852. DOI: 10.1109/TBME.2009.2035102.

(50) Cheng, Y. H.; Chen, Y. C.; Brien, R.; Yoon, E. Scaling and automation of a high-throughput single-cell-derived tumor sphere assay chip. Lab Chip 2016, 16 (19), 3708–3717. DOI: 10.1039/c6lc00778c.

(51) Chen, Y. C.; Zhang, Z.; Yoon, E. Early Prediction of Single-Cell Derived Sphere Formation Rate Using Convolutional Neural Network Image Analysis. Anal Chem 2020, 92 (11), 7717–7724. DOI: 10.1021/acs.analchem.0c00710.

(52) Hartnett, E. B.; Zhou, M.; Gong, Y. N.; Chen, Y. C. LANCE: a Label-Free Live Apoptotic and Necrotic Cell Explorer Using Convolutional Neural Network Image Analysis. Anal Chem 2022, 94 (43), 14827–14834. DOI: 10.1021/acs.analchem.2c00878.

(53) Love, M. I.; Huber, W.; Anders, S. Moderated estimation of fold change and dispersion for RNA-seq data with DESeq2. Genome Biol 2014, 15 (12), 550. DOI: 10.1186/s13059-014-0550-8 From NLM Medline.

(54) Langmead, B.; Salzberg, S. L. Fast gapped-read alignment with Bowtie 2. Nat Methods 2012, 9 (4), 357–359. DOI: 10.1038/nmeth.1923 From NLM Medline.

(55) Subramanian, A.; Tamayo, P.; Mootha, V. K.; Mukherjee, S.; Ebert, B. L.; Gillette, M. A.; Paulovich, A.; Pomeroy, S. L.; Golub, T. R.; Lander, E. S.;, et al. Gene set enrichment analysis: a knowledge-based approach for interpreting genome-wide expression profiles. Proc Natl Acad Sci U S A 2005, 102 (43), 15545–15550. DOI: 10.1073/pnas.0506580102.

(56) Liberzon, A.; Subramanian, A.; Pinchback, R.; Thorvaldsdottir, H.; Tamayo, P.; Mesirov, J. P. Molecular signatures database (MSigDB) 3.0. Bioinformatics 2011, 27 (12), 1739–1740. DOI: 10.1093/bioinformatics/btr260.

(57) Shannon, P.; Markiel, A.; Ozier, O.; Baliga, N. S.; Wang, J. T.; Ramage, D.; Amin, N.; Schwikowski, B.; Ideker, T. Cytoscape: a software environment for integrated models of biomolecular interaction networks. Genome Res 2003, 13 (11), 2498–2504. DOI: 10.1101/gr.1239303.

(58) Guha, R. Chemical informatics functionality in R. Journal of Statistical Software 2007, 18, 1–16.

(59) Kim, S.; Thiessen, P. A.; Cheng, T.; Yu, B.; Bolton, E. E. An update on PUG-REST: RESTful interface for programmatic access to PubChem. Nucleic Acids Res 2018, 46 (W1), W563–W570. DOI: 10.1093/nar/gky294 From NLM Medline.

(60) Kim, S.; Thiessen, P. A.; Bolton, E. E.; Bryant, S. H. PUG-SOAP and PUG-REST: web services for programmatic access to chemical information in PubChem. Nucleic Acids Res 2015, 43 (W1), W605–611. DOI: 10.1093/nar/gkv396 From NLM Medline.

(61) Gartel, A. L. FOXM1 in Cancer: Interactions and Vulnerabilities. Cancer Res 2017, 77 (12), 3135–3139. DOI: 10.1158/0008-5472.CAN-16-3566.

(62) Katzenellenbogen, B. S.; Guillen, V. S.; Katzenellenbogen, J. A. Targeting the oncogenic transcription factor FOXM1 to improve outcomes in all subtypes of breast cancer. Breast Cancer Res 2023, 25 (1), 76. DOI: 10.1186/s13058-023-01675-8.

(63) Balsas, P.; Galan-Malo, P.; Marzo, I.; Naval, J. Bortezomib resistance in a myeloma cell line is associated to PSMbeta5 overexpression and polyploidy. Leuk Res 2012, 36 (2), 212–218. DOI: 10.1016/j.leukres.2011.09.011.

(64) Manasanch, E. E.; Orlowski, R. Z. Proteasome inhibitors in cancer therapy. Nature Reviews Clinical Oncology 2017, 14 (7), 417–433. DOI: 10.1038/nrclinonc.2016.206.

(65) Tew, K. D. Commentary on “Proteasome Inhibitors: A Novel Class of Potent and Effective Antitumor Agents”. Cancer Research 2016, 76 (17), 4916–4917. DOI: 10.1158/0008-5472.Can-16-1974.

(66) Qiu, Z.; Oleinick, N. L.; Zhang, J. ATR/CHK1 inhibitors and cancer therapy. Radiother Oncol 2018, 126 (3), 450–464. DOI: 10.1016/j.radonc.2017.09.043.

(67) Dent, P.; Tang, Y.; Yacoub, A.; Dai, Y.; Fisher, P. B.; Grant, S. CHK1 inhibitors in combination chemotherapy: thinking beyond the cell cycle. Mol Interv 2011, 11 (2), 133–140. DOI: 10.1124/mi.11.2.11.

(68) Nolan, T. J.; Lok, J. B. Macrocyclic lactones in the treatment and control of parasitism in small companion animals. Curr Pharm Biotechnol 2012, 13 (6), 1078–1094. DOI: 10.2174/138920112800399167.

(69) Merola, V. M.; Eubig, P. A. Toxicology of avermectins and milbemycins (macrocylic lactones) and the role of P-glycoprotein in dogs and cats. Vet Clin North Am Small Anim Pract 2012, 42 (2), 313–333, vii. DOI: 10.1016/j.cvsm.2011.12.005.

(70) Avcioglu, H.; Balkaya, I. A comparison of the efficacy of subcutaneously administered ivermectin, doramectin, and moxidectin against naturally infected Toxocara vitulorum in calves. Trop Anim Health Prod 2011, 43 (6), 1097–1099. DOI: 10.1007/s11250-011-9807-3.

(71) Ballweber, L. R.; Smith, L. L.; Stuedemann, J. A.; Yazwinski, T. A.; Skogerboe, T. L. The effectiveness of a single treatment with doramectin or ivermectin in the control of gastrointestinal nematodes in grazing yearling stocker cattle. Vet Parasitol 1997, 72 (1), 53–68. DOI: 10.1016/s0304-4017(97)00078-2.

(72) Chen, C.; Liang, H.; Qin, R.; Li, X.; Wang, L.; Du, S.; Chen, Z.; Meng, X.; Lv, Z.; Wang, Q.;, et al. Doramectin inhibits glioblastoma cell survival via regulation of autophagy in vitro and in vivo. Int J Oncol 2022, 60 (3). DOI: 10.3892/ijo.2022.5319.

(73) Chu, W. Y.; Dorlo, T. P. C. Pyronaridine: a review of its clinical pharmacology in the treatment of malaria. J Antimicrob Chemother 2023, 78 (10), 2406–2418. DOI: 10.1093/jac/dkad260.

(74) Bailly, C. Pyronaridine: An update of its pharmacological activities and mechanisms of action. Biopolymers 2021, 112 (4), e23398. DOI: 10.1002/bip.23398.

(75) Puhl, A. C.; Gomes, G. F.; Damasceno, S.; Godoy, A. S.; Noske, G. D.; Nakamura, A. M.; Gawriljuk, V. O.; Fernandes, R. S.; Monakhova, N.; Riabova, O.;, et al. Pyronaridine Protects against SARS-CoV-2 Infection in Mouse. ACS Infect Dis 2022, 8 (6), 1147–1160. DOI: 10.1021/acsinfecdis.2c00091.

(76) Lane, T. R.; Massey, C.; Comer, J. E.; Freiberg, A. N.; Zhou, H.; Dyall, J.; Holbrook, M. R.; Anantpadma, M.; Davey, R. A.; Madrid, P. B.;, et al. Pyronaridine tetraphosphate efficacy against Ebola virus infection in guinea pig. Antiviral Res 2020, 181, 104863. DOI: 10.1016/j.antiviral.2020.104863.

(77) Villanueva, P. J.; Martinez, A.; Baca, S. T.; DeJesus, R. E.; Larragoity, M.; Contreras, L.; Gutierrez, D. A.; Varela-Ramirez, A.; Aguilera, R. J. Pyronaridine exerts potent cytotoxicity on human breast and hematological cancer cells through induction of apoptosis. PLoS One 2018, 13 (11), e0206467. DOI: 10.1371/journal.pone.0206467.

(78) Villanueva, P. J.; Gutierrez, D. A.; Contreras, L.; Parra, K.; Segura-Cabrera, A.; Varela-Ramirez, A.; Aguilera, R. J. The Antimalarial Drug Pyronaridine Inhibits Topoisomerase II in Breast Cancer Cells and Hinders Tumor Progression In Vivo. Clin Cancer Drugs 2021, 8 (1), 50–56. DOI: 10.2174/2212697X08666210219101023.

(79) Kobayashi, S.; Shimamura, T.; Monti, S.; Steidl, U.; Hetherington, C. J.; Lowell, A. M.; Golub, T.; Meyerson, M.; Tenen, D. G.; Shapiro, G. I.;, et al. Transcriptional profiling identifies cyclin D1 as a critical downstream effector of mutant epidermal growth factor receptor signaling. Cancer Res 2006, 66 (23), 11389–11398. DOI: 10.1158/0008-5472.CAN-06-2318.

(80) Zhong, Z. H.; Yi, Z. L.; Zhao, Y. D.; Wang, J.; Jiang, Z. B.; Xu, C.; Xie, Y. J.; He, Q. D.; Tong, Z. Y.; Yao, X. J.;, et al. Pyronaridine induces apoptosis in non-small cell lung cancer cells by upregulating death receptor 5 expression and inhibiting epidermal growth factor receptor. Chem Biol Drug Des 2022, 99 (1), 83–91. DOI: 10.1111/cbdd.13926 From NLM Medline.

(81) Zhu, C.; Wei, Y.; Wei, X. AXL receptor tyrosine kinase as a promising anti-cancer approach: functions, molecular mechanisms and clinical applications. Mol Cancer 2019, 18 (1), 153. DOI: 10.1186/s12943-019-1090-3.

(82) Scaltriti, M.; Elkabets, M.; Baselga, J. Molecular Pathways: AXL, a Membrane Receptor Mediator of Resistance to Therapy. Clin Cancer Res 2016, 22 (6), 1313–1317. DOI: 10.1158/1078-0432.CCR-15-1458.

(83) Zhou, M.; Ma, Y.; Rock, E. C.; Chiang, C. C.; Luker, K. E.; Luker, G. D.; Chen, Y. C. Microfluidic single-cell migration chip reveals insights into the impact of extracellular matrices on cell movement. Lab Chip 2023, 23 (21), 4619–4635. DOI: 10.1039/d3lc00651d.

(84) Tanaka, M.; Siemann, D. W. Therapeutic Targeting of the Gas6/Axl Signaling Pathway in Cancer. Int J Mol Sci 2021, 22 (18). DOI: 10.3390/ijms22189953 From NLM Medline.

(85) Tanaka, M.; Siemann, D. W. Gas6/Axl Signaling Pathway in the Tumor Immune Microenvironment. Cancers (Basel) 2020, 12 (7). DOI: 10.3390/cancers12071850 From NLM PubMed-not-MEDLINE.

(86) Zhang, Y.; Arner, E. N.; Rizvi, A.; Toombs, J. E.; Huang, H.; Warner, S. L.; Foulks, J. M.; Brekken, R. A. AXL Inhibitor TP-0903 Reduces Metastasis and Therapy Resistance in Pancreatic Cancer. Mol Cancer Ther 2022, 21 (1), 38–47. DOI: 10.1158/1535-7163.MCT-21-0293.

(87) Shen, Y.; Chen, X.; He, J.; Liao, D.; Zu, X. Axl inhibitors as novel cancer therapeutic agents. Life Sci 2018, 198, 99–111. DOI: 10.1016/j.lfs.2018.02.033.

(88) Costello, J. C.; Heiser, L. M.; Georgii, E.; Gonen, M.; Menden, M. P.; Wang, N. J.; Bansal, M.; Ammad-ud-din, M.; Hintsanen, P.; Khan, S. A.;, et al. A community effort to assess and improve drug sensitivity prediction algorithms. Nat Biotechnol 2014, 32 (12), 1202–1212. DOI: 10.1038/nbt.2877.

(89) Chiu, Y. C.; Zheng, S.; Wang, L. J.; Iskra, B. S.; Rao, M. K.; Houghton, P. J.; Huang, Y.; Chen, Y. Predicting and characterizing a cancer dependency map of tumors with deep learning. Sci Adv 2021, 7 (34). DOI: 10.1126/sciadv.abh1275 From NLM Medline.

(90) Shi, X.; Gekas, C.; Verduzco, D.; Petiwala, S.; Jeffries, C.; Lu, C.; Murphy, E.; Anton, T.; Vo, A. H.; Xiao, Z.;, et al. Building a translational cancer dependency map for The Cancer Genome Atlas. Nat Cancer 2024, 5 (8), 1176–1194. DOI: 10.1038/s43018-024-00789-y.

(91) Corsello, S. M.; Nagari, R. T.; Spangler, R. D.; Rossen, J.; Kocak, M.; Bryan, J. G.; Humeidi, R.; Peck, D.; Wu, X.; Tang, A. A.;, et al. Discovering the anti-cancer potential of non-oncology drugs by systematic viability profiling. Nat Cancer 2020, 1 (2), 235–248. DOI: 10.1038/s43018-019-0018-6.

(92) Singh, N. R.; Morris, C. M.; Koleth, M.; Wong, K.; Ward, C. M.; Stevenson, W. S. Polyploidy in myelofibrosis: analysis by cytogenetic and SNP array indicates association with advancing disease. Mol Cytogenet 2013, 6 (1), 59. DOI: 10.1186/1755-8166-6-59.

(93) Chen, E.; Beer, P. A.; Godfrey, A. L.; Ortmann, C. A.; Li, J.; Costa-Pereira, A. P.; Ingle, C. E.; Dermitzakis, E. T.; Campbell, P. J.; Green, A. R. Distinct clinical phenotypes associated with JAK2V617F reflect differential STAT1 signaling. Cancer Cell 2010, 18 (5), 524–535. DOI: 10.1016/j.ccr.2010.10.013.

(94) Zachos, G.; Black, E. J.; Walker, M.; Scott, M. T.; Vagnarelli, P.; Earnshaw, W. C.; Gillespie, D. A. Chk1 is required for spindle checkpoint function. Dev Cell 2007, 12 (2), 247–260. DOI: 10.1016/j.devcel.2007.01.003.

(95) Yu, X.; Li, W.; Liu, H.; Deng, Q.; Wang, X.; Hu, H.; Xu-Monette, Z. Y.; Xiong, W.; Lu, Z.; Young, K. H.;, et al. Ubiquitination of the DNA-damage checkpoint kinase CHK1 by TRAF4 is required for CHK1 activation. J Hematol Oncol 2020, 13 (1), 40. DOI: 10.1186/s13045-020-00869-3.

(96) Herrtwich, L.; Nanda, I.; Evangelou, K.; Nikolova, T.; Horn, V.; Sagar Erny, D.; Stefanowski, J.; Rogell, L.; Klein, C.;, et al. DNA Damage Signaling Instructs Polyploid Macrophage Fate in Granulomas. Cell 2016, 167 (5), 1264–1280 e1218. DOI: 10.1016/j.cell.2016.09.054.

(97) Ullah, Z.; de Renty, C.; DePamphilis, M. L. Checkpoint kinase 1 prevents cell cycle exit linked to terminal cell differentiation. Mol Cell Biol 2011, 31 (19), 4129–4143. DOI: 10.1128/MCB.05723-11.

